# Identification of drug modifiers for RYR1 related myopathy using a multi-species discovery pipeline

**DOI:** 10.1101/813097

**Authors:** Jonathan Volpatti, Yukari Endo, Linda Groom, Stephanie Brennan, Ramil Noche, William Zuercher, Peter Roy, Robert T. Dirksen, James J. Dowling

## Abstract

Ryanodine receptor type I-related myopathies (RYR1-RMs) represent the largest group of non-dystrophic myopathies. RYR1-RMs are associated with severe disabilities and early mortality; despite these facts, there are currently no available treatments. The goal of this study was to identify new therapeutic targets for RYR1-RMs. To accomplish this, we developed a novel discovery pipeline using nematode, zebrafish, and mammalian cell models of the disease. We first performed large-scale drug screens in *C. elegans* and zebrafish. 74 positive hits were identified in *C. elegans*, while none were uncovered in the zebrafish. Targeted testing of these hits in zebrafish yielded positive results for two compounds. We examined these compounds using newly created *Ryr1* knockout C2C12 cells, and found that p38 inhibition impaired caffeine-induced Ca^2+^ release. Lastly, we tested one p38 inhibitor in myotubes from *Ryr1^Y524S/+^* (YS) mice, and demonstrated that it blunts the aberrant temperature-dependent increase in resting Ca^2+^ in these cells. In all, we developed a unique platform for RYR1-RM therapy development that is potentially applicable to a broad range of neuromuscular disorders.

## Introduction

The ryanodine receptor type I (RyR1) is a calcium release channel located in the terminal cisternae of the sarcoplasmic reticulum (SR) in skeletal muscle. During excitation-contraction coupling (ECC), RyR1 is activated by the voltage sensing L-type calcium channel dihydropyridine receptor (DHPR), located in the transverse tubule (T-tubule) membrane. Together, the T-tubule and two adjacent SR terminal cisternae form a junctional membrane unit referred to as the triad (Jungbluth, 2007; Dowling et al., 2014; Jungbluth et al. 2018). Mutations in the *RYR1* gene are the most common cause of non-dystrophic muscle disease in humans (Colombo et al., 2015; Gonorazky, et al.; 2018; Jungbluth et al., 2018). *RYR1* mutations are associated with a wide range of clinical phenotypes, collectively referred to as RYR1-related myopathies (RYR1-RM), that can include wheelchair and ventilator dependence, and dynamic symptoms such as exercise induced myalgias, heat stroke, and malignant hyperthermia (Klein et al., 2012; Amburgey et al., 2013; Snoeck, et al., 2015; Jungbluth et al., 2016; Matthews et al., 2018). Despite their relatively high prevalence and associated morbidities, there are currently no approved pharmacological therapies for patients with RYR1-RM.

Much of what is known about the function of RyR1 and the impact of its mutations on skeletal muscle come from animal models. Well described recessive models of RYR1-RM include the *C. elegans unc-68* mutant (null mutant with impaired motility (Maryon et al., 1996; Maryon et al., 1998), the *relatively relaxed* zebrafish (loss of function *ryr1b* mutant with impaired motility and early death [Hirata et al., 2007]), and the “dyspedic” *Ryr1* null mouse (perinatal lethal [Buck et al., 1997; Avila and Dirksen, 2000]). In addition, two compound heterozygous mouse models of recessive RYR1-RM were with recently generated and characterized (Brennan et al., 2019; Elbaz et al., 2019). These models are complimented by “knock-in” mutants in mice that mirror specific dominant human mutations, including the I4895T mutant (associated with central core disease and referred to as the IT model) (Zvaritch et al., 2007; Zvaritch et al., 2009; Lee et al., 2017), the R163C mutant (associated with malignant hyperthermia) (Yang et al., 2006), and the Y522S mutant (associated with malignant hyperthermia and referred to as the YS mouse) (Chelu et al., 2006; Durham et al., 2008; Lanner et al., 2012; Yarotskyy et al., 2013).

Previous work using these models identified potential therapeutic targets for RYR1-RM (for a comprehensive review, see Lawal et al., 2018) including anti-oxidants (Durham et al., 2008; Dowling et al., 2012; Michelucci et al., 2017), ER stress modulators (Lee et al., 2017), and chemicals that influence the binding of RyR1 to modifying partners (e.g. S107, which promotes RyR1/calstabin1 interaction) (Lehnart et al., 2008; Bellinger et al., 2008; Andersson et al., 2011). However, as of yet none of these targets has successfully translated to patients, though *N*-acetylcysteine was tested in a recently completed clinical trial (ClinicalTrials.gov identifier: NCT02362425), where it failed to achieve its primary endpoint (Todd et al., *Neurology*, in press). There is thus a critical need to identify and develop new treatment strategies.

With the goal of identifying new therapies for RYR1-RM, we set out to establish a novel multi-species translational pipeline (Figure 1). This pipeline is based on the functional conservation of RyR1 across many species, and takes advantage of specific attributes of *C. elegans* (ability to rapidly screen thousands of compounds), zebrafish (large scale testing in a vertebrate model), and mammalian cell lines (translatability to humans). We screened several thousand compounds, and identified a p38 inhibitor that modified RyR1 phenotypes in all three systems. Our study identifies a new potential therapeutic strategy for RYR1-RM, outlines the utility of multi-species drug discovery, and lays the groundwork for future similar screens for other neuromuscular disorders.

**Figure 1.**
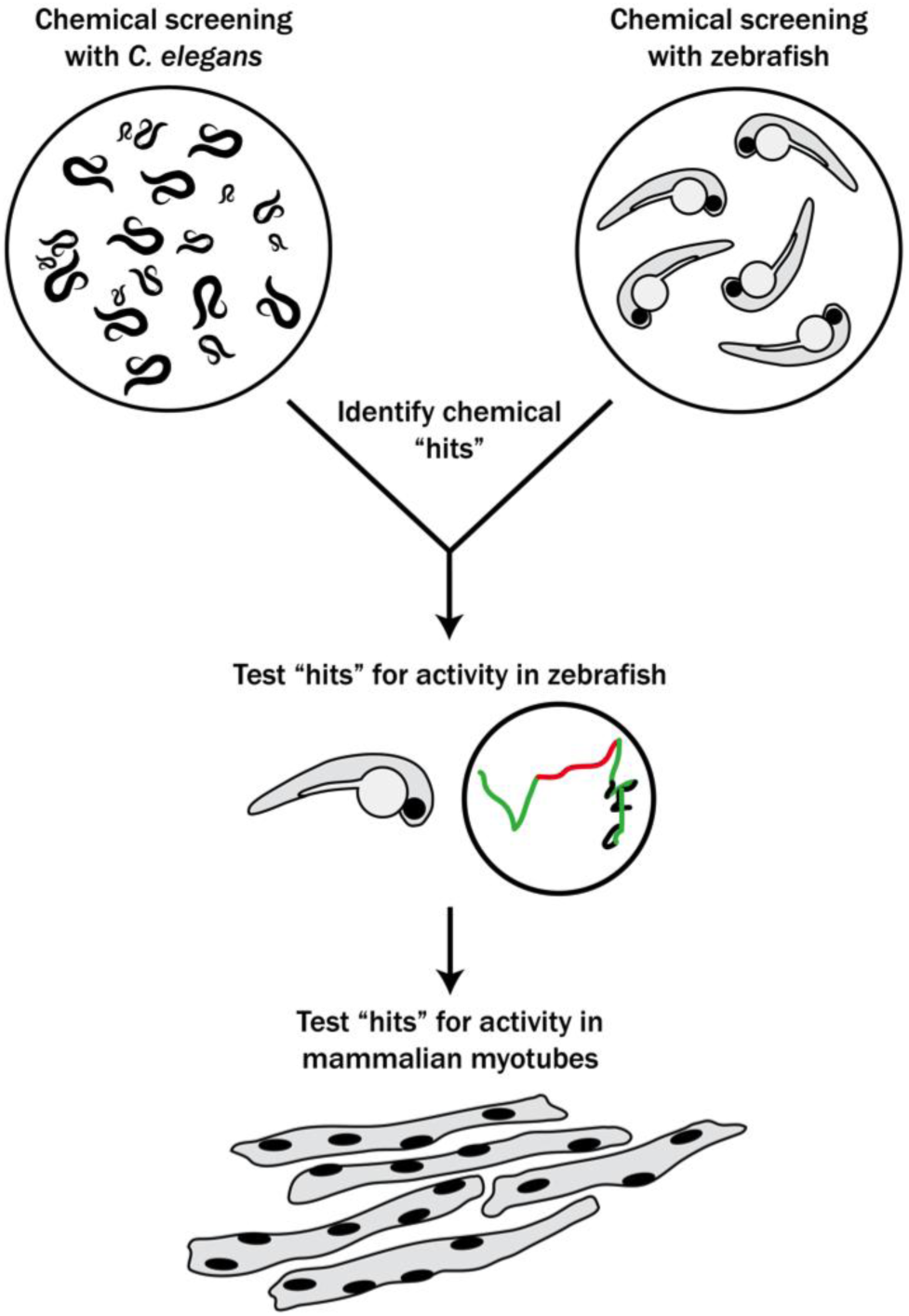
Schematic of our multi-species translational pipeline aimed at identifying potential therapeutic targets for RYR1-RM. The pipeline involved screening *C. elegans* and zebrafish with thousands of compounds for suppressors of RYR1 mutant phenotypes, followed by further characterization in zebrafish and evaluation in mammalian cell lines.

## Results

### Large scale chemical screen in C. elegans identifies 74 unc-68 suppressors

We performed a drug screen using the *unc-68 C. elegans* model of RYR1-RM (Figure 2). This model has a loss of expression mutation in the worm ryanodine receptor (Maryon et al., 1998). The *unc-68* mutation results in a characteristic “uncoordinated” (*unc*) movement phenotype, impaired calcium regulation, and reduced survival. We considered using a liquid-based movement assay (i.e., the *C. elegans* “thrashing assay” [Maryon et al., 1996; Maryon et al., 1998]) as the basis for our drug screen because *unc-68* mutants have been reported to thrash at lower rates than WT (see Supplementary Figure 1A), but found that automatable methods for this assay were sufficiently variable to prevent use in a screen of thousands of compounds. Instead, we developed a sensitized screen based on our observation that nemadipine-A, an inhibitor of the dihydropyridine receptor (Kwok et al., 2006), induces developmental growth arrest in *unc-68* mutants (Supplementary Figure 1B). Specifically, *unc-68* worms exposed to 25 µM nemadipine arrest at the L1-L3 larval stage, while the majority of wild type N2 strain treated with nemadipine have either normal development or, in a small percentage, arrest at the L4 stage (Supplementary Figure 1B). Based on this, we screened for chemicals that could overcome this growth arrest.

**Figure 2.**
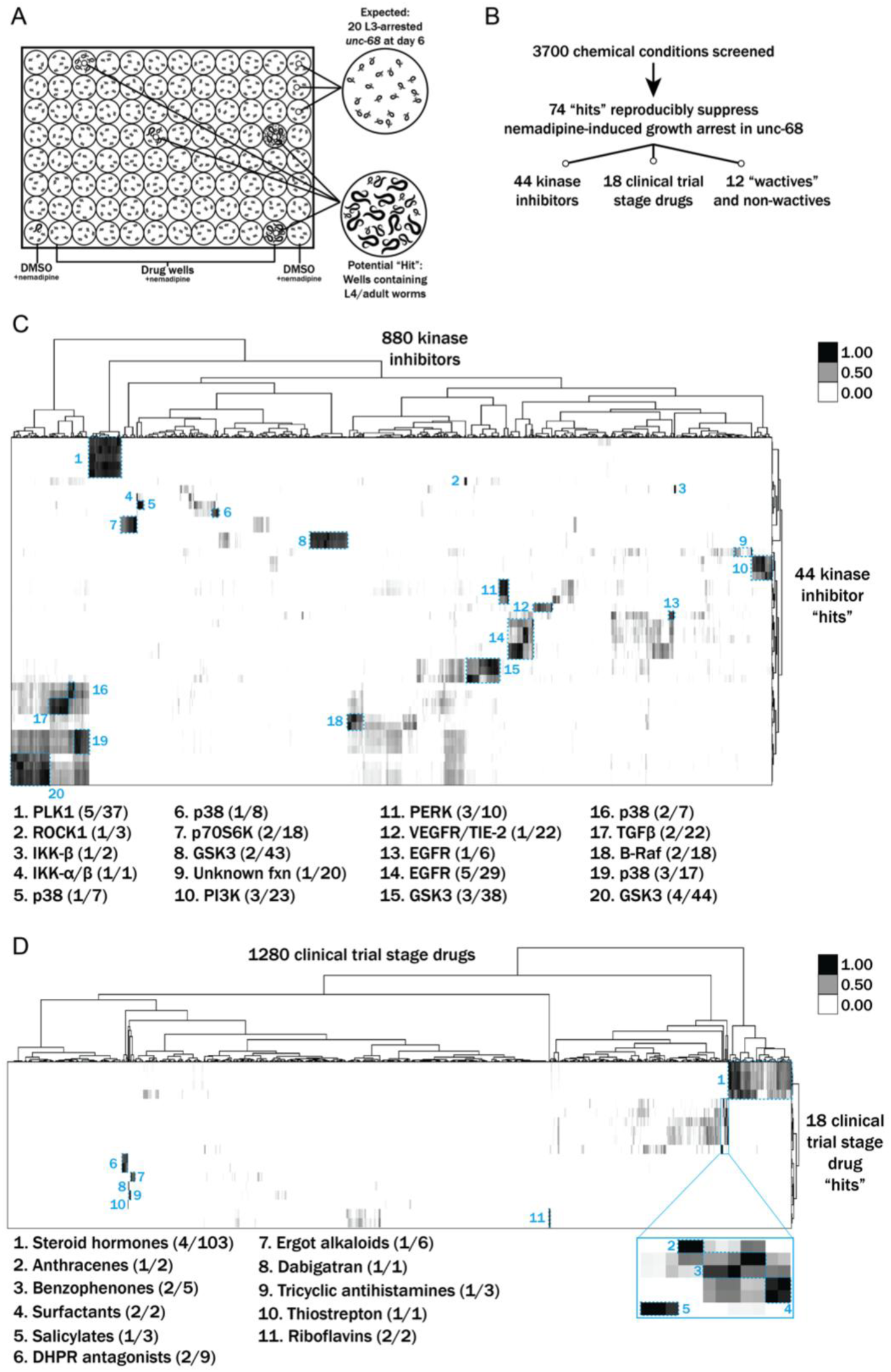
Results from a chemical suppressor screen of *unc-68* mutants. **A)** Schematic of our screen methodology showing the expected growth arrest phenotype of *unc-68* worms exposed to 25 μM nemadipine after 6 days of exposure and the expected phenotype of a chemical that suppresses this growth arrest. **B)** Summary of the 74 “hits” from this screen that reproducibly suppressed nemadipine-induced growth arrest of *unc-68* mutants. **C-D)** Heat map visualization of hits from the screen. All of the compounds screened from each library are arranged on the x-axis, while “hits” are arranged on the y-axis. Tanimoto scores were calculated for each pair of compounds as a measure of structural similarity and similar clusters were identified via hierarchical clustering of Tanimoto scores. As shown, chemicals with similar structures are associated with similar annotated functions/targets. Fisher’s exact test was used to determine enrichment based on the number of structurally similar members in each cluster that were either hits or not hits out of the total number of the compounds in the library.

We evaluated 3700 conditions from a combination of libraries: 770 <u>w</u>orm-bio<u>active</u> (a.k.a. “wactives”) and non-wactives at 60 µM and 7.5 µM (Burns et al., 2015), 1280 drugs from the US Drug Collection at 60 µM (MicroSource Discovery Systems), and 880 proprietary kinase inhibitors at 60 µM. We found 278 chemicals overcame the developmental arrest induced by nemadipine in duplicate wells (Figure 2B). However, 62 single wells with DMSO as a control also contained a few *unc-68* mutants that reached the L4 stage; thus, we concluded that some of the 278 chemicals were likely false positives. We prioritized and re-tested 145 chemicals that strongly suppressed the phenotype in duplicate and found 74 chemical “hits” which reproducibly suppressed the nemadipine-induced growth suppression of *unc-68* mutants (Figure 2B). Among these 74 “hits”, we identified five wactives and seven non-wactives, 44 kinase inhibitors, and 18 compounds from the MicroSource library (Supplementary Table I).

Many of the chemicals that were screened have redundant targets and/or functions (e.g. there are several EGFR inhibitors in the kinase library and several steroid hormones in the US Drug Collection). Therefore, we reasoned that hits that are overrepresented among groups of structurally and/or functionally related chemicals contained within the libraries ought to be prioritized for further testing in zebrafish. To determine enrichment based on structural similarity, we compared the chemical structures of the hits with those of the chemicals in each library by hierarchical clustering of their Tanimoto scores (Figure 2C-D) (Burns et al., 2015). This method shows how structurally similar chemicals cluster with one another.

Using this methodology, p38 inhibitors were revealed to be significantly overrepresented (\*\**p*=0.0022) among the 44 positive hits from the kinase inhibitor library. Interestingly, structurally dissimilar p38 inhibitors suppressed the phenotype (clusters 5, 6, 16, and 19 in Figure 2C), suggesting that inhibition of their common target is responsible for the activity. Additionally, we found that EGFR, PERK, and PLK1 inhibitors were overrepresented (\*\**p*=0.0060, \**p*=0.0109, \**p*=0.0326, respectively). Lastly, we identified multiple GSK3 (n=9) and PI3K (n=3) inhibitors that strongly suppressed the phenotype, but these were not statistically overrepresented among the hits (*p*=0.4093 and *p*=0.1027, respectively).

Among the 18 positive hits from the MicroSource library, we found several classes of chemical that were overrepresented (Figure 2D). These included riboflavins (\*\*\**p*=0.0002), surfactants (\*\*\**p*=0.0002), benzophenones (\*\**p*=0.0018), anthracenes (\**p*=0.0279), salicylates (\**p*=0.0416), and tricyclic antihistamines (\**p*=0.0416). Interestingly, two DHPR inhibitors out of nine present in the library (\*\**p*=0.0063) were identified as suppressors of the growth arrest induced by nemadipine, itself a DHPR inhibitor. This may reflect competition for receptor binding which diminishes the effect of nemadipine, or perhaps a higher effective concentration which becomes agonistic to the receptor. Of note, riboflavin and riboflavin 5-phosphate sodium were both identified as hits and they appear to be structurally distinct from every other chemical in the US Drug Collection (cluster 11, Figure 2D). Similarly, thiostrepton and dabigatran etexilate mesylate are both structurally unique molecules and overrepresented among the 18 hits (\**p*=0.0141).

Altogether, the overrepresented groups of chemicals and structurally unique molecules among the “hits” were prioritized for follow-up testing in *ryr1b* mutant zebrafish.

### Large scale chemical screen in ryr1 zebrafish

In parallel, we performed a screen in a zebrafish model of RYR1-RM. Zebrafish have two *ryr1* paralogs. Recessive mutations in *ryr1a* cause no overt phenotype, while recessive mutations in *ryr1b* result in abnormal swim behavior and early lethality after 11-13 days of life (Hirata et al., 2007). *ryr1a*; *ryr1b* double mutants exhibit no movement and have a median survival of 5 days of life (Supplementary Figure 2A and 2B; Chagovetz et al., 2019). We used the double mutants for our screen because of their obvious motor phenotype and because variability in the *ryr1b* single mutant motor phenotype precluded large-scale screening (Supplementary Figure 2C). We screened 436 kinase inhibitors from the DiscoveryProbe Kinase Inhibitor Library (ApexBio) and 1360 drugs from the US Drug Collection (MicroSource Discovery Systems) for improvement in motility of the *ryr1a*; *ryr1b* double mutants (as measured by touch-evoked escape response) (Figure 3). We did not identify a single compound that was able to promote movement in these fish.

**Figure 3.**
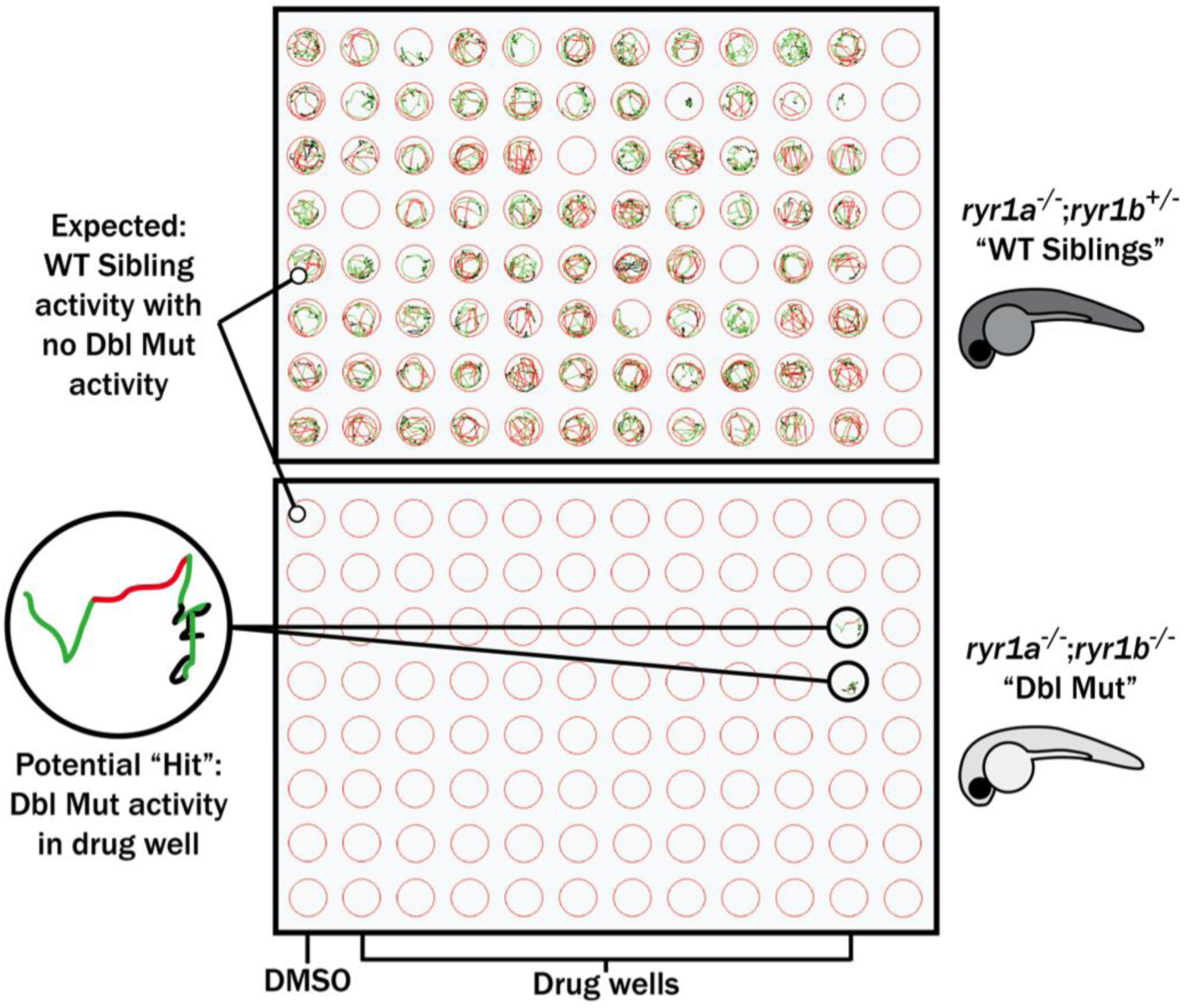
Schematic of our zebrafish screen methodology showing the expected motility of unaffected siblings (“WT Siblings”) and double mutants (“Dbl Mut”) and expected motility of immotile double mutants if a chemical suppressed the phenotype. We did not identify any suppressors of the double mutant phenotype.

### Targeted drug testing in ryr1 zebrafish

Using our prioritized list based on enrichment modeling, we next sought to determine if any positive hits from our *C. elegans* screen could improve phenotypes in either the *ryr1a*; *ryr1b* double mutants or the *ryr1b* mutant zebrafish. We used motility as our outcome measure, examining both touch-evoked escape response and optogenetically induced swimming as previously utilized by our lab (Sabha et al., 2016). Among the overall group of 74 hits, we tested 4/5 wactives, 7/7 non-wactives, 16/18 MicroSource drugs, and several inhibitors from the six classes of kinase inhibitors over-represented among the hits.

First, we tested all of these at a single concentration of 10 µM for the ability to promote movement in the *ryr1a*; *ryr1b* double mutants, which in the untreated state lack any movement. Consistent with our large scale screen, none of these chemicals improved the double mutant phenotype. We then tested all of these chemicals for the ability to suppress the abnormal touch-evoke escape response of *ryr1b* mutants, a transient phenotype that is most apparent between 3-4 days of life (Hirata et al., 2007). The chemical hits did not modulate this phenotype either.

Lastly, we assayed many of these chemicals on the *ryr1b* mutants to determine their effect on movement speed using the optogenetic motility assay. We analyzed for a positive chemical-genetic interaction, which we defined as an unexpected movement phenotype not resulting from the combined additive or multiplicative effects of genotype and chemical treatment. We found positive chemical-genetic interactions for p38 and PI3K inhibitors (Figure 4A/B). For several of these compounds, the effect was characterized by a worsening of WT motility due to the chemical whereas the *ryr1b* mutants appear unaffected by the chemical. Additionally, we found positive interactions with two wactives (Figure 4C). We did not see any positive chemical-genetic interplay with other “hits” from the MicroSource library; in addition, some of the MicroSource compounds were toxic at the tested concentrations. In all, the molecules that promote defects in WT but not *ryr1* mutant zebrafish likely represent chemicals that act as negative modifiers of signaling pathways related to RyR1, or act directly on RyR1, but in its absence have no effect.

**Figure 4.**
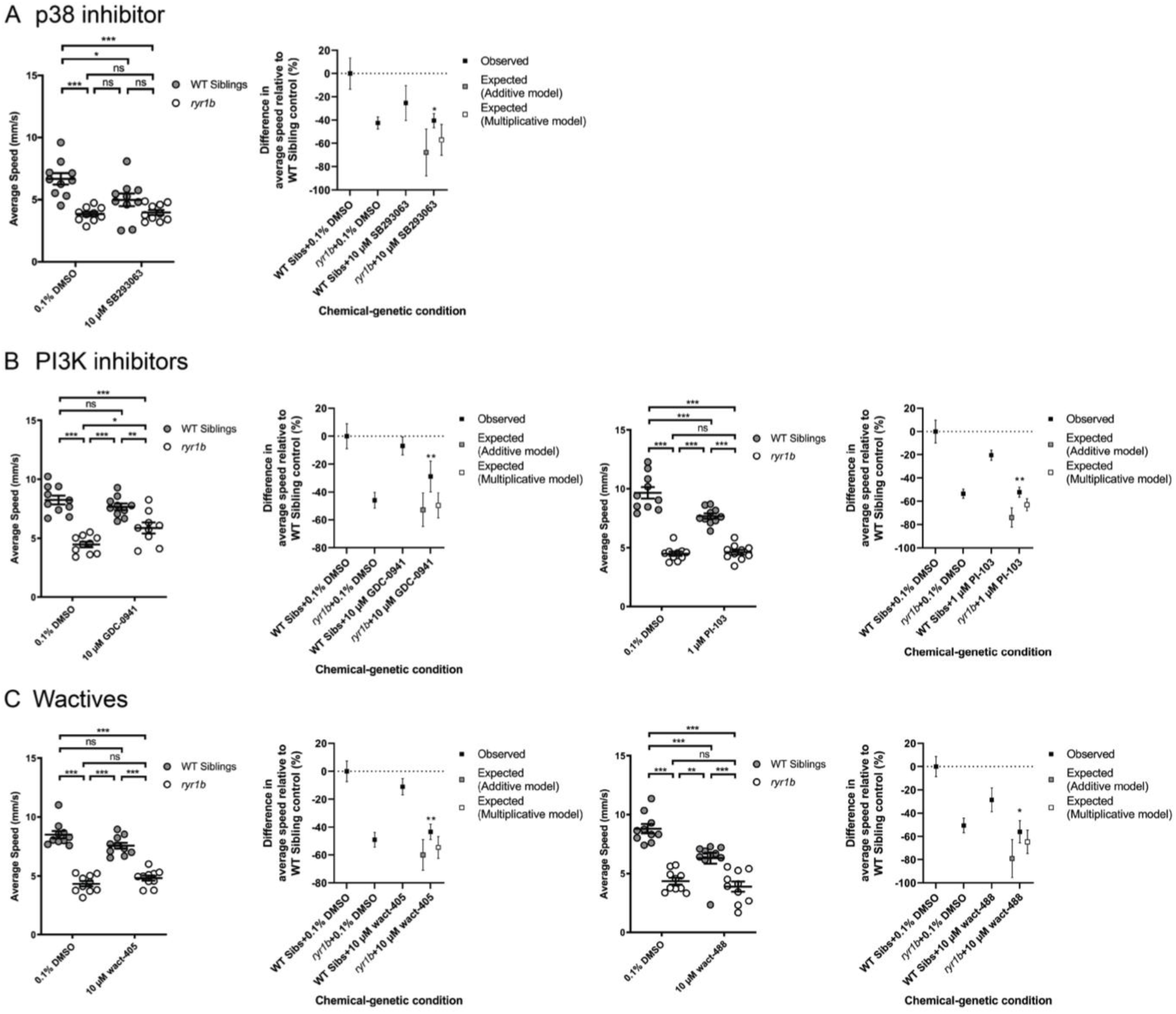
Positive chemical-genetic effects of **A)** a p38 inhibitor, **B)** two PI3K inhibitors, and **C)** two “wactives” on *ryr1b* mutant movement speed. In each set, the average speed of individual larvae is shown on the left, while the difference compared to WT controls (i.e. WT siblings+DMSO vehicle) is given on the right. Statistical significance was determine by two-way ANOVA with Tukey’s multiple comparison post-test. For the difference compared to WT controls, significance indicates an interaction between genotype and chemical treatment such that the combination contributes a large percentage of total variation in the data, greater or less than what would be expected by an additive or multiplicative model.

### Testing positive hits in C2C12 myotubes

We lastly sought to examine the potential translatability of our findings to mammalian models of RYR1-RM. To accomplish this, we tested the effect of two positive hits, identified in *C. elegans* and *ryr1b* mutant zebrafish (p38 inhibitor SB202190 and PI3K inhibitor GDC-0941), on RyR1-dependent Ca^2+^ release in C2C12 mouse myotubes. We examined this in wild type C2C12s and in a new generated *Ryr1* knockout line. We created this line using CRISPR/Cas9 gene editing; it contains a bi-allelic frameshift deletion mutation in *Ryr1* (which we refer to as “KO”). Successful targeting of the *Ryr1* locus was demonstrated by Sanger sequencing, lack of off target mutation verified by whole genome sequencing, and absence of RyR1 protein expression confirmed by Western blot analysis (Supplementary Figure 3).

We measured intracellular calcium release from RyR1 in response to acute application of 10 mM caffeine (Tong et al., 1997; Meissner, 2017) in control and *Ryr1* KO C2C12 myotubes after 24 hours incubation with either SB202190 or GDC-0941. As expected, untreated *Ryr1* KO myotubes lacked caffeine-induced Ca^2+^ release; this not change upon incubation with 1 µM GDC-0941(Supplementary Figure 4). Interestingly, and consistent with our data in zebrafish, SB202190 impaired caffeine-induced calcium release in control WT C2C12 myotubes (Figure 5A, C). Given this finding, we tested an additional p38 inhibitor, SB203580, on WT cells and observed a dose dependent inhibition of calcium release (Figure 5B). These data thus suggest that p38 inhibition disrupts calcium release from RyR1, identifying p38 inhibitors as novel modulators of RyR1 signalling. Of note, both p38 inhibitors promoted essentially no change in caffeine-induced calcium release in KO cells, with the exception of a very small increase with 100 nM SB202190 that was not seen with 10 uM.

**Figure 5.**
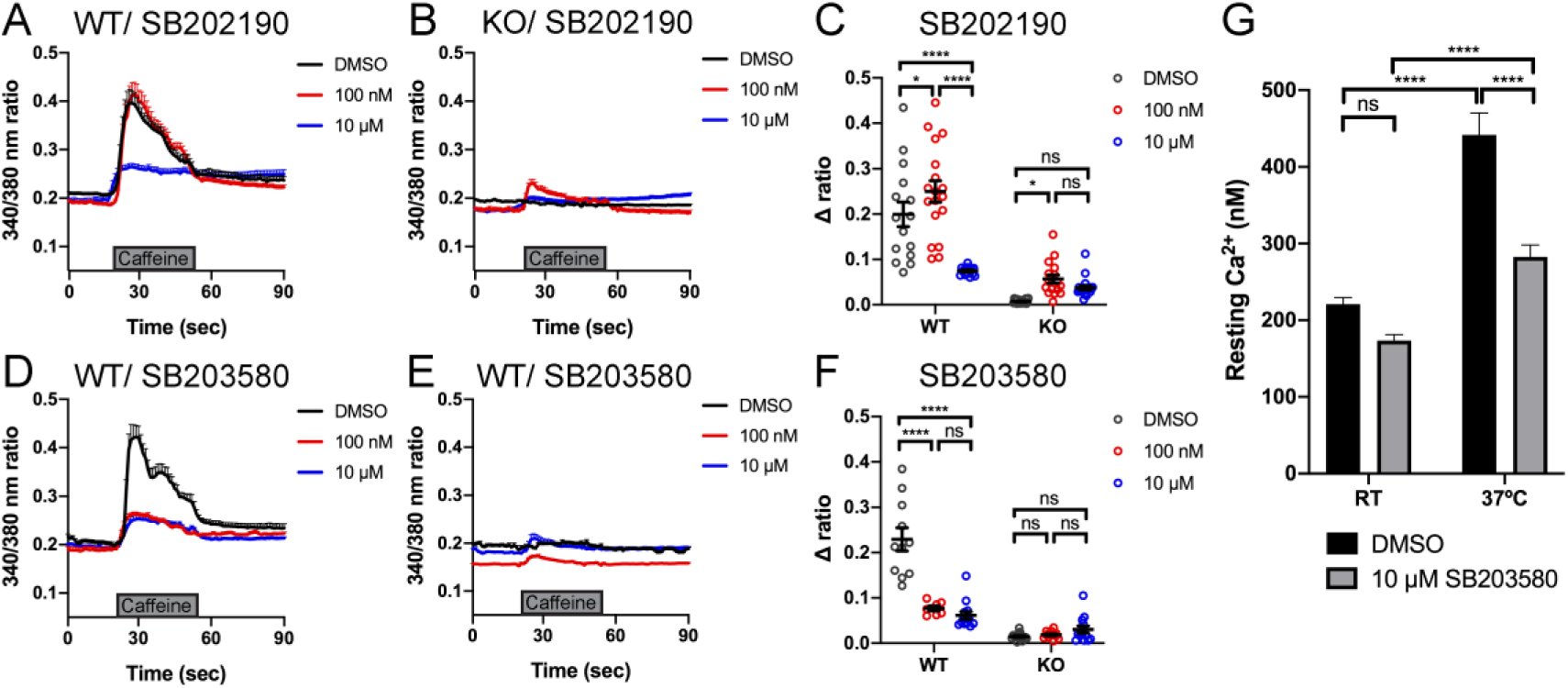
Intracellular calcium measurement in myotubes. **A-B)** Ratiometric fura-2 imaging with 10 mM caffeine induction after SB202190 treatment. **A)** WT: DMSO; n=15, 100 nM; n=18, 10 μM; n=14. **B)** KO: DMSO; n=19, 100 nM; n=17, 10 μM; n=22. Data are presented as mean ± SEM. **C)** Peak changes in intracellular Ca^2+^ of (A) and (B) expressed as Δratio (Ratio peak – Ratio baseline). WT: DMSO vs 100 nM; \**p* = 0.0486, DMSO vs 10 μM; \*\*\*\**p* < 0.0001, 100 nM vs 10 μM; \*\*\*\**p* < 0.0001. KO: DMSO vs 100 nM; \**p* = 0.0406, DMSO vs 10 μM; p = 0.2564, 100 nM vs 10 μM; p = 0.5730. **D-E)** Ratiometric fura-2 imaging with 10 mM caffeine induction after SB203580 treatment. **D)** WT: DMSO; n=11, 100 nM; n=7, 10 μM; n=13. **E)** KO: DMSO; n=15, 100 nM; n=12, 10 μM; n=13. Data are presented as mean ± SEM. Statistical analysis by two-way ANOVA followed by Tukey’s multiple comparisons post-test. **F)** Peak changes in intracellular Ca^2+^ of (C) and (D) expressed as Δratio (Ratio peak – Ratio baseline). WT: DMSO vs 100 nM; \*\*\*\**p* < 0.0001, DMSO vs 10 μM; \*\*\*\**p* < 0.0001, 100 nM vs 10 μM; p = 0.6663. KO: DMSO vs 100 nM; p = 0.9661, DMSO vs 10 μM; p = 0.5441, 100 nM vs 10 μM; p = 0.7296. Statistical analysis by two-way ANOVA followed by Tukey’s multiple comparisons post-test. **G)** Resting cytosolic Ca^2+^ in YS myotubes. At room temperature (RT); with DMSO; n = 24, with 10 μM SB203580; n = 31. At 37°C; with DMSO; n = 24, with 10 μM SB203580; n = 31. RT DMSO vs RT SB203580; p = 0.2355, RT DMSO vs 37°C DMSO; \*\*\*\**p* < 0.0001, RT SB203580 vs 37°C SB203580; \*\*\*\**p* < 0.0001, 37°C DMSO vs 37°C SB203580; \*\*\*\**p*<0.0001. Statistical analysis by two-way ANOVA followed by Sidak’s multiple comparisons post-test.

### SB203580 abrogates the temperature-dependent increase in resting Ca^2+^ in YS myotubes

The fact that p38 inhibitors negatively modulate RyR1 calcium release in WT zebrafish and C2C12 cells opens the possibility that they may serve as modulators of phenotypes related to RyR1 hyperexcitability. To test this possibility, we examined calcium dynamics in myotubes from the YS mouse model of RYR1-RM. The YS model contains a point mutation analogous to the Y522S mutation found in patients with malignant hyperthermia and central core pathology (Quane et al., 1994). The YS mutation enhances both the sensitivity of RyR1 to activators (e.g. DHPR, caffeine, 4-chloro-*m*-cresol) and the temperature-dependence of RyR1 Ca^2+^ leak, alterations that underlie the MH susceptibility and exertional heat stroke phenotypes of these mice (Chelu et al., 2006; Durham et al., 2008; Lanner et al., 2012). We examined if the temperature dependent increase in resting myoplasmic Ca^2+^ concentration in YS myotubes was abrogated by the p38 inhibitor SB203580. Overnight incubation of YS myotubes with 10 µM SB203580 significantly reduced the temperature-dependent increase in resting Ca^2+^ observed in YS myotubes (Figure 5C).

### Discussion

In this study, we developed a novel multi-system pipeline for drug discovery and development for RYR1-RM. Using this platform, we were able to screen several thousand compounds and test their efficacy across multiple species. We identified p38 inhibitors as a new class of potential modifiers of RyR1 signalling. This platform can be applied to new compounds that may improve RYR1-RM phenotypes, as well as for additional diseases with suitable animal models.

The strengths of our pipeline include the rapidity with which we are able to screen drugs and the potential for increased translatability in drugs that positively modify a diverse group of *in vivo* models. In terms of speed, the initial screen in *C. elegans* was completed within two weeks, while the large-scale screen in zebrafish was accomplished in one month. While this does not approach the speed and scale of cell culture screens, it is fast and efficient and importantly is done *in vivo* using outcome measures that are relatable to human disease phenotypes. It would additionally be possible to add a cell culture based screen into the pipeline as a pre-screen or in parallel with the animal model testing. For example, studies for RyR1 functional interactors are underway using novel calcium indicators such as SERCaMP (Henderson et al., 2014); these indicators may prove ideal for screens in disease relevant cell models. Recently, a high-throughput screen was performed with HEK293 cells that successfully identified novel inhibitors of RyR1 (Murayama et al., 2018), supporting the potential utility of a cell-based screen in identifying modifiers of mutant RyR1 activity.

In terms of translatability, it is difficult to say whether a multi-organism strategy is superior to other approaches and/or more likely to yield targets that will work in humans. This is because, at present, no drugs have successfully been translated to patients with RYR1-RM. However, there is reason to speculate that a compound that can modify phenotype(s) associated with dysfunction of a gene in these diverse evolutionary settings may promote improvement in a more universal way.

One of the primary shortcomings of our study is the relative lack of positive hits, particularly in the zebrafish. We believe this is primarily related to the specific nature of our existing zebrafish model, which completely lacks expression of RyR1. There are currently no human patients with bi-allelic null mutations, and thus our fish models do not accurately mirror the human disease. Furthermore, it is likely that no compounds can compensate for the complete loss of RyR1 activity in terms of live animal phenotypes. One of our primary future directions is therefore to develop new models of RYR1-RM that are better suited to drug discovery (such as in the worm and in the fish) and for subsequent testing in mammals. In this vein, there are very recent publications detailing new recessive RYR1-RM mouse models including a p.G2435R mutant (Lopez et al., 2018), a p.A4329D/p.Q1970fsX16 compound heterozygote model (Elbaz et al., 2019), and a p.T4706M/indel compound heterozygote model generated by our group (Brennan et al., 2019). In addition, seven RYR1-RM equivalent mutations in *unc-68* were modeled by transgenic overexpression in *C. elegans* and these mutants exhibited hypersensitivity to caffeine and the malignant hyperthermia triggering agent halothane (Baines et al., 2017).

As the primary purpose of this study was to develop a new drug screening platform and test its potential efficacy, we did not focus further on mechanism(s) of action of any of our positive hits. It would, of course, be of interest to understand how p38 inhibition is serving to modify RyR1 dependent calcium release and related phenotypes. One hypothesis is that targets downstream of p38 impair calcium entry into ER stores. Alternatively, the inhibitors may be modifying the phosphorylation state of RyR1, which in turn may impact channel function. Interestingly, p38α/β MAPK activity is elevated in aged skeletal muscle stem cells (i.e. satellite cells), and inhibition of p38 helps to restore the regenerative capacity of these stem cells (Cosgrove et al., 2014). This raises the intriguing possibility that p38 inhibition modulates a Ca^2+^-dependent pathway in both skeletal muscle and closely associated satellite cells. Most relevant to future clinical intervention of RYR1-RM, it was shown recently that combined treatment of *N*-acetylcysteine and the p38 inhibitor SB203580 lead to robust expansion of satellite cell populations *in vitro* and *in vivo* (L’honoré et al., 2018). Future work, beyond the scope of the current study, will be required to establish drug mechanisms, and to test the potential for polytherapy.

### Conclusion

In all, we established a unique “multi species” pipeline for drug discovery for RYR1-RM. This platform is rapid and robust, and provides the ability to examine multiple different types of *in vivo* models. Our study lays the ground work for its future use in RYR1-RM drug development, and for establishment of similarly platforms for other rare diseases.

## Materials and Methods

### Animal ethics statement

All zebrafish experiments were performed in accordance with all relevant ethical regulations, specifically following the policies and guidelines of the Canadian Council on Animal Care and an institutionally reviewed and approved animal use protocol (#41617). No additional ethical approval was required for our experiments with the invertebrate nematode worm *C. elegans*.

### Chemical sources

Chemical libraries used in this study include the US Drug Collection (1280 compounds; MicroSource Discovery Systems Inc.), the DiscoveryProbe Kinase Inhibitor Library (436 compounts; APExBIO), a proprietary kinase inhibitor library (880 compounds), and a collection of 770 <u>w</u>orm-bio<u>actives</u> (“wactives”) that were identified in a screen for bioactive small molecules in *C. elegans* (Burns et al., 2015). Nemadipine-A (#5619779), optovin analog 6b8 (#5707191), and wactives/non-wactives (ID numbers in Supplementary Table I) were purchased from ChemBridge. Chemicals from the US Drug Collection identified as “hits” in the *C. elegans* screen were purchased from Sigma-Aldrich for testing in zebrafish. Kinase inhibitors representative of those identified as “hits” in the *C. elegans* screen, including p38 inhibitors, were purchased individually from APExBIO or selected from the DiscoveryProbe Kinase Inhibitor Library for testing in zebrafish and cell lines.

### C. elegans strains and culture

All animals were cultured under standard methods at 20 °C (Burns et al., 2015). The wildtype (N2) and *unc-68* (TR2170 and TR2171) strains of *Caenorhabditis elegans* were obtained from the C. elegans Genetics Center (University of Minnesota). The “thrashing assay” is performed by counting the waveforms propagated by individual *C. elegans* in one minute as previously described (Maryon et al., 1996).

### C. elegans chemical screening

The protocol for the 96-well liquid-based chemical screens was described previously (Burns et al., 2015). Briefly, nematode growth media (NGM; for recipe see Burns et al., 2015) was used to concentrate saturated *Escherichia coli* culture (strain HB101) two-fold (NGM-HB101). Nemadipine A (NEM) was added to NGM+HB101 to a final concentration of 31.25 µM/0.5% DMSO (NEM+NGM+HB101). A total of 40 µL of NEM+NGM+HB101 was dispensed into each well of a 96-well plate, and 300 nL of chemical dissolved in DMSO was pinned into the wells using a 96-well pinning tool (V&P Scientific). At Day 0, approximately 20 synchronized first larval-stage (L1), *unc-68* (TR2171) worms obtained from an embryo preparation were added to each well in 10 µL of M9 buffer (Burns et al., 2015). The final concentration of dimethyl sulfoxide (DMSO) in the wells was 1% v/v. Plates were sealed with parafilm, wrapped in damp paper towels to reduce evaporation in wells, and incubated for 6 days at 20°C while shaking at 200 rpm (New Brunswick I26/I26R shaker, Eppendorf). A stereomicroscope was used to assess developmental stage after 6 days incubation. All preliminary screens and re-tests were performed in duplicate.

### Cheminformatics

Pairwise Tanimoto coefficient scores were calculated for each chemical “hit” and compound in the screening library using OpenBabel (http://openbabel.org) as previously described (Burns et al., 2015). Tanimoto coefficient scores were hierarchically clustered in Cluster 3.0 using an unweighted Euclidean distance similarity metric with complete linkage clustering and visualized in TreeView as previously described (Folts et al., 2016). For enrichment analysis, chemicals were counted based on the number of structurally similar members in each cluster (Tanimoto scores were >0.55 for the majority of members in a given cluster) and Fisher’s exact test (GraphPad Prism 8) was used to calculate enrichment.

### Zebrafish care and husbandry

In this study, we used *ryr1a* and *ryr1b* mutant alleles that have been previously characterized (Hirata et al., 2007; Chagovetz et al., 2019). Both mutants result in loss of RYR1 protein expression from the mutant allele. For follow-up screens, we generated single *ryr1b*^-/-^ mutants via incross of *ryr1b*^+/-^ carriers. Phenotypic analysis of all *ryr1* mutants was performed on a stereomicroscope.

### Chemical treatments

All chemical stocks were prepared in DMSO and added to egg water at 0.1%-0.5% of the final volume to prepare working concentrations (depending on chemical solubility). Equal volumes of vehicle solvent were used in all conditions for a single assay. Note that methylene blue was not added to the egg water. Dishes or 96-well plates were sealed with parafilm, wrapped in aluminum foil, and incubated at 28.5°C until the assay date. Different volumes and culture dish formats were used depending on the endpoint assay.

### Zebrafish chemical screening

The US Drug Collection (1280 compounds) and the DiscoveryProbe Kinase Inhibitor Library (436 compounts) were screened for chemicals that could promote motility in *ryr1a; ryr1b* double mutants. Library stocks of 10 mM in DMSO were added to 0.1% of the final volume to egg water to prepare a 10 µM screening concentration. Specifically, 150 µL of egg water was added to every well of two separate 96-well plates. Double mutants were generated by in-cross of *ryr1a*^-/-^;*ryr1b*^+/-^ mutants. Embryos were manually dechorionated at 1 dpf. Then at 2 dpf two double mutant embryos were added to each well of one plate while two phenotypically wildtype siblings were added to the second plate as a control. Care was taken to lower embryos to the bottom of a Pasteur pipette tip and deposit embryos into the well by surface tension to minimize changes to well volume. Next, 250 µL of the drug library was prepared at 40 µM working concentration in a separate 96-well plate by adding 1 µL of the 10 mM stock to 249 µL of egg water. Next, 50 µL of the 40 µM working concentration was added to the 150 µL water containing embryos to give a final concentration of 10 µM drug. After 24 hours incubation in chemical, motility of 3 dpf wildtype and double mutant larvae was assessed by touch-evoked response (Hirata et al., 2007) under a stereomicroscope.

### Photoactivation of motor behaviour assay

At 3 dpf, *ryr1b* mutants were segregated from wild-type (*ryr1b*^+/+^ or *ryr1b*^+/-^) siblings based on their phenotype and distributed into sterile 6 cm tissue culture dishes containing 10 mL of egg water plus chemical. All assays were performed after 24 hour incubation at 4 dpf using ZebraBox platform (ViewPoint) and 10 µM optovin analog 6b8 (ID 5707191; ChemBridge) as previously described (Sabha et al., 2016).

### Generating Ryr1 knockout C2C12 cells using CRISPR-Cas9 strategy

The original C2C12 (ATCC® CRL1772™) was purchased from American Type Culture Collection (Manassas, VA, USA). The sgRNA sequence (5’-AGGAGAGAAGGTTCGAGTTG-3’) against *Ryr1* in Exon 6 was designed by the online CRISPR Design Tool (http://tools.genome-engineering.org) and cloned at the BbsI site into pSpCas9(BB)-2A-Puro (PX459) V2.0 (Addgene plasmid ID: 62988). The CRISPR plasmids are transfected into C2C12 cells by electroporation (Amaxa Nucleofector^TM^ Lonza). Seventy-two hours later, the transfected cells were selected in medium containing 4 µg/mL puromycin for three days and then subcloned into 96-well plates. Once at sufficient cell density, the genomic DNA of subclones was analyzed by sanger sequencing. Primers used were (forward) 5’-GTGTGACGGGAGTCCCAAAT-3’ and (reverse)5’-ACTGGGCATGCCAATGATGA-3’.

### Western blot

Protein was isolated in RIPA buffer from C2C12 myotubes at 5days after starting differentiation. Cells are incubated with 100 nM SB202190, 10 µM SB203580, DMSO and without treatment for 24 hours from day4. A total of 30 µg of total protein was run on either 4.5% SDS-acrylyamide gel for RyR1 or 15% SDS-acrylamide gel for β-actin. Blots were run for 3 – 3.5 h for Ryr1 or for 2 – 2.5 h for βactin at 100 V and transferred overnight at 20 V. The membrane was blocked in 3% bovine serum albumin (BSA) in Tris Buffered Saline with Tween 20 (TBST) for 1 h at room temperature before incubating with primary antibodies overnight at 4°C. Antibodies used were anti-Ryanodine receptor antibody 34C (Developmental Studies Hybridoma Bank) at 1:100 dilution and anti-beta actin antibody (mAbcam 8226, abcam) at 1:5000 dilution. After three washes in TBST, blots were incubated with Anti-Mouse IgG-HRP conjugate (Bio-Rad) at 1:10000 dilution. Blots were imaged by chemiluminescence (Western Lightning Plus-ECL, PerkinElmer) using the Gel Doc™ XR + Gel Documentation System (BioRad), and band signal intensities determined using ImageLab software (BioRad).

### Intracellular Ca^2+^ measurements in myotubes

Intracellular Ca^2+^ measurements were obtained from Fura-2 AM-loaded myotubes as described previously (Goonasekera et al., 2007). Briefly, myotubes grown on glass bottom dishes were loaded with 5 µM Fura-2 AM for 45 min at 37°C in a normal rodent Ringer’s solution consisting of (in mM): 145 NaCl, 5 KCl, 2 CaCl_2_, 1 MgCl_2_, 10 HEPES, pH 7.4. Coverslips of Fura-2– loaded cells were then mounted in a tissue chamber on the stage of an epifluorescence-equipped inverted microscope (Zeiss). Cells were sequentially excited at 340- and 380-nm wavelength and fluorescence emission at 510 nM was collected using a high-speed CCD camera (Hamamatsu). The results are presented as the ratio of F340/F380. Maximal increase in intracellular Ca^2+^ by induction of 10 mM caffeine was defined as the difference between peak and baseline fluorescence ratios.

### Statistical analysis

Tests of statistical significance were performed using Microsoft Office Excel 2008 (Microsoft) and GraphPad Prism 8 for Mac OSX (GraphPad Software). Differences were considered to be statistically significant at *P* less than 0.05 (*), *P* < 0.01 (**), or *P* < 0.001 (***). All data unless otherwise specified are presented as mean ± SEM.

## Acknowledgements

The authors wish to thank Dr. Andrew Burns at University of Toronto for helpful discussion and insight. We wish to thank Dr. David Grunwald at University of Massachusetts Medical School for generously providing the *ryr1a* zebrafish mutants. The study was funded by grants from Muscular Dystrophy Association and the RYR1 Foundation (JJD and RTD).

---

**Supplementary Figure 1.**
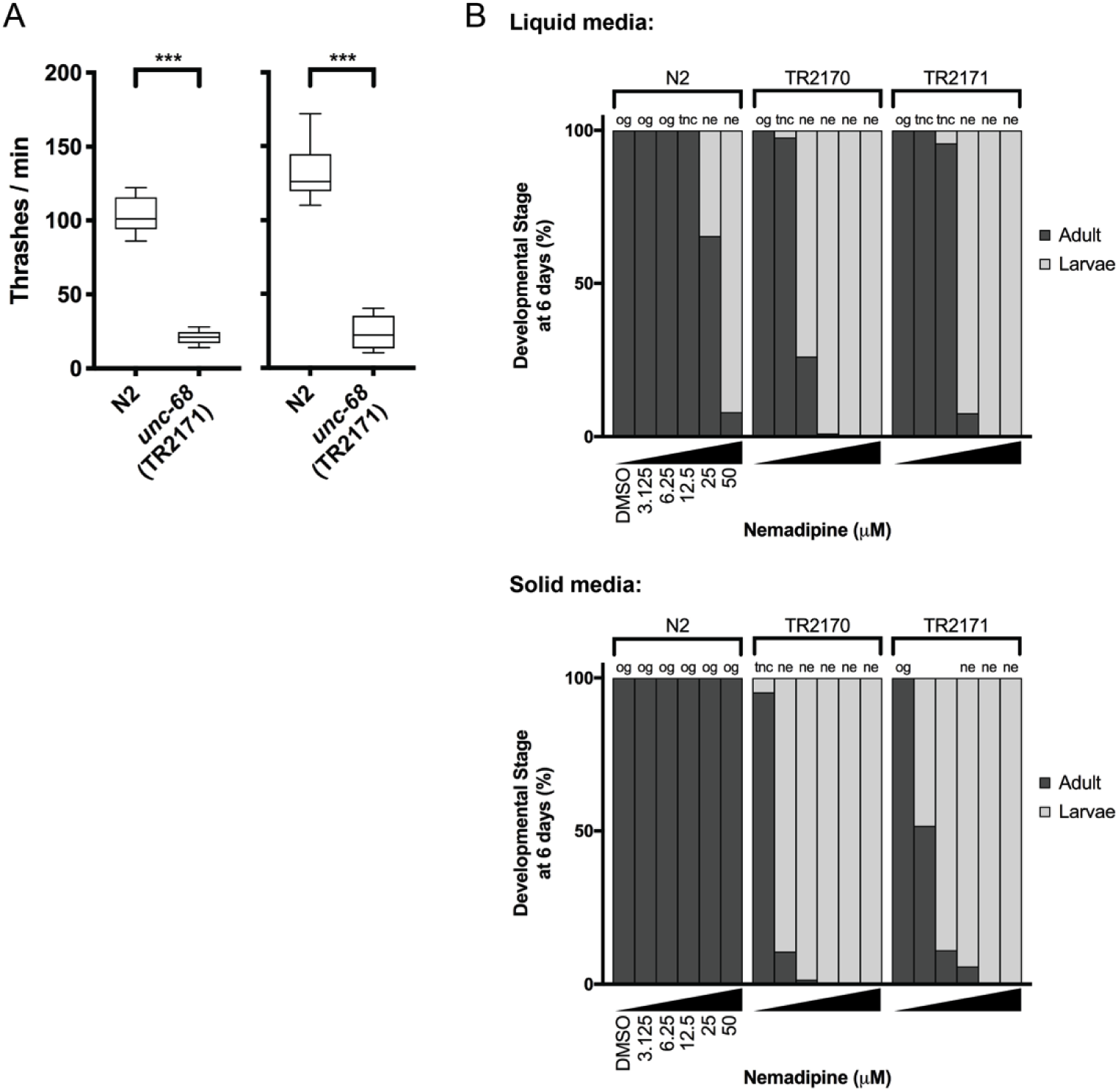
**A)** The “thrashing assay” is performed by counting the waveforms propagated by individual *C. elegans* in one minute (Maryon et al., 1996). Consistent with previous reports (Maryon et al., 1996; Maryon et al., 1998), we found that *unc-68* mutants thrash at a significantly lower rate than WT in two independent assays (*n* = 10 per genotype; Student’s T-test: \*\*\**P*<0.001). **B)** In both liquid and solid media, the DHPR inhibitor nemadipine-A (Kwok et al., 2006) induces concentration-dependent developmental growth arrest in *unc-68* mutants. L1 larvae were plated in chemical at Day 0 and developmental stage was measured at Day 6. Abbreviations: og = overgrown with larvae/eggs and food source depleted; tnc = larvae and eggs were too numerous to count, but food source not depleted; ne: no eggs have been laid (this phenotype is consistent with inhibition of egg-laying defective protein EGL-19, a DHPR, by nemadipine-A).

**Supplementary Figure 2.**
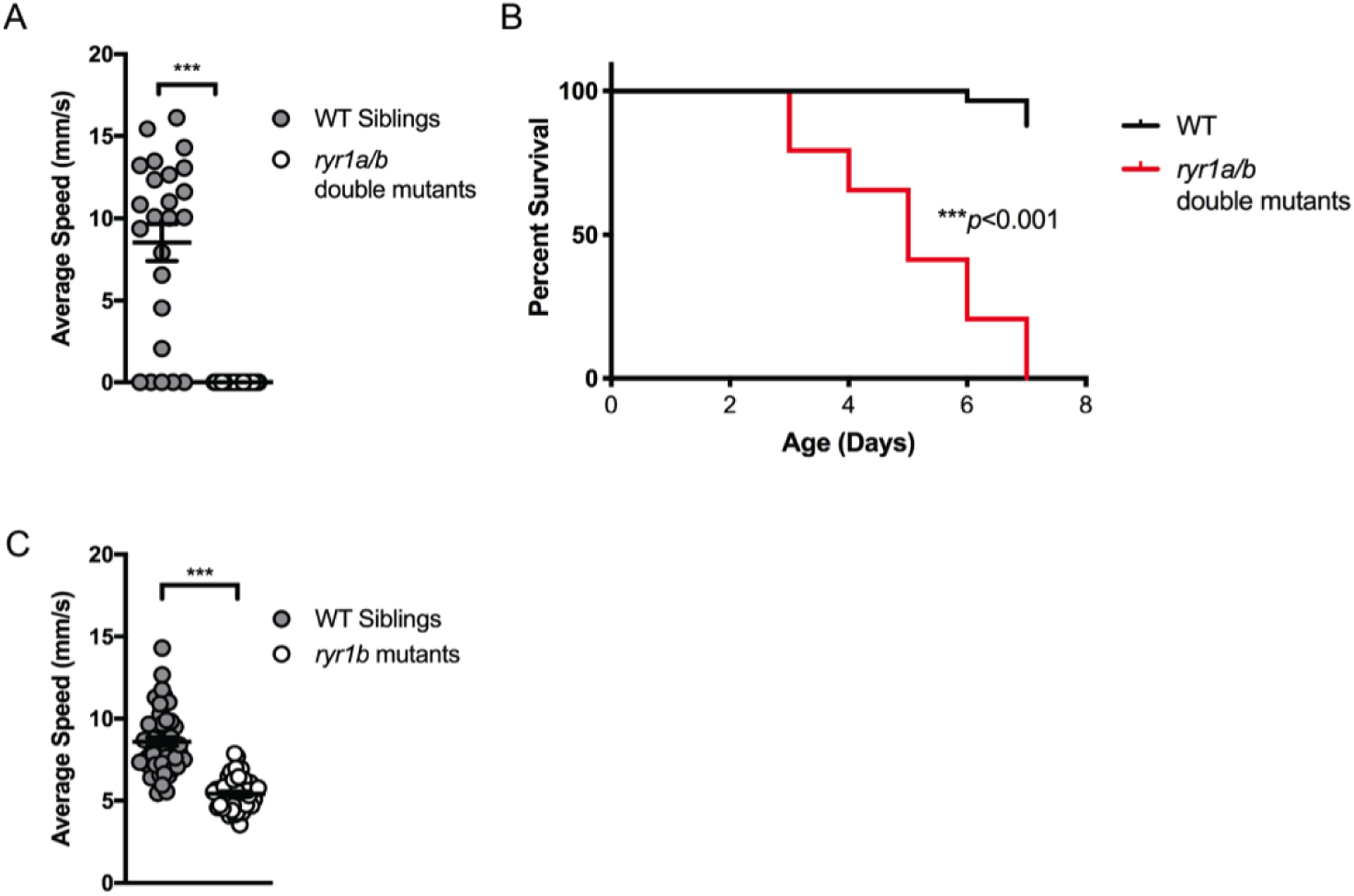
**A)** *ryr1a*;*ryr1b* double mutants exhibit complete lack of movement as shown by optovin-6b8 induced swimming. They do not respond to touch stimuli either (data not shown; independently shown by Chagovetz et al., 2019). **B)** The *ryr1a*;*ryr1b* double mutants exhibit early larval lethality with a median survival of 5 days and maximum survival of 7 days. **C)** The ryr1b mutants show significantly reduced movement speed at 4 dpf by optovin-6b8 induced swimming. However, there is sufficient overlap with WT siblings to preclude large-scale screening.

**Supplementary Figure 3.**
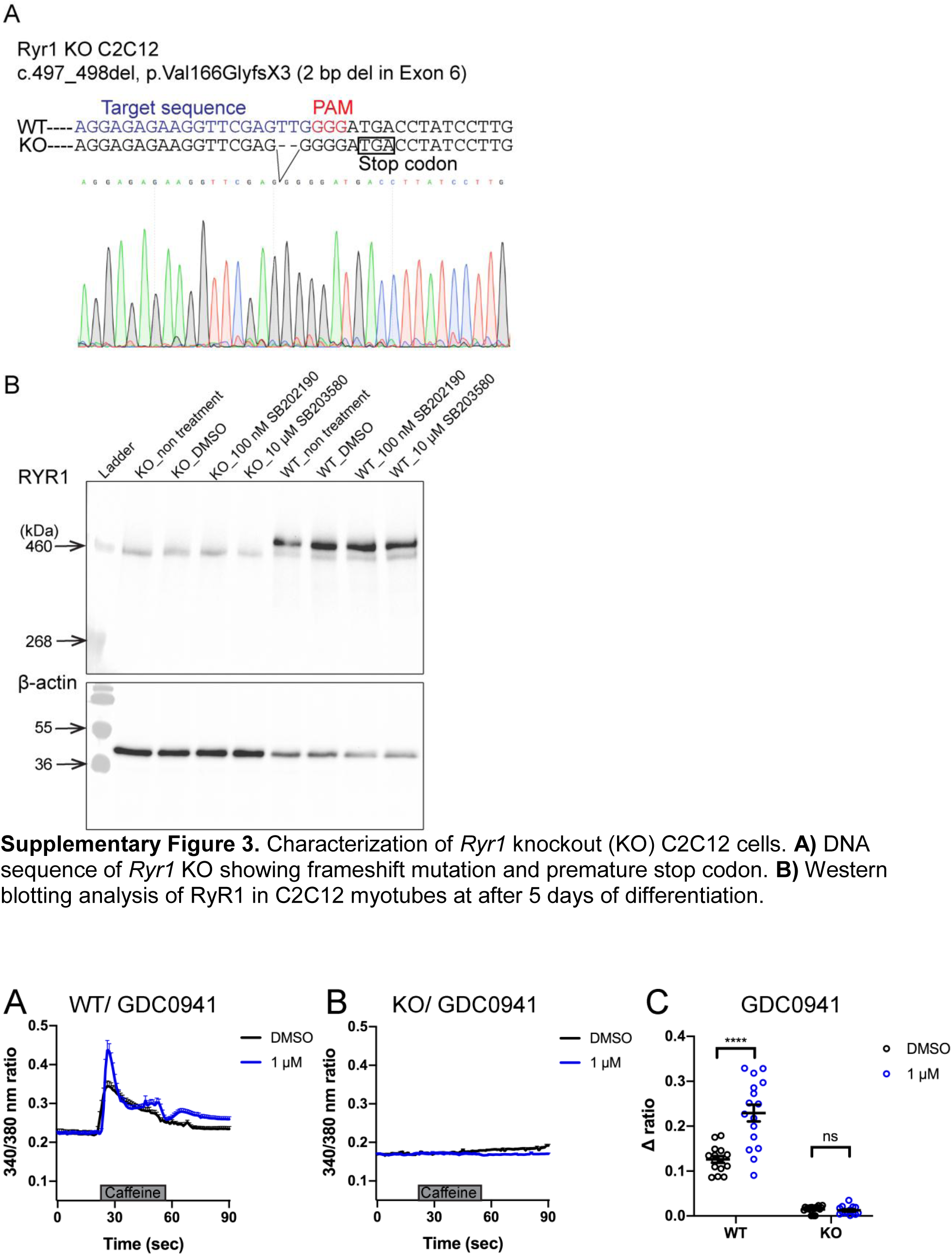
Characterization of *Ryr1* knockout (KO) C2C12 cells. **A)** DNA sequence of *Ryr1* KO showing frameshift mutation and premature stop codon. **B)** Western blotting analysis of RyR1 in C2C12 myotubes at after 5 days of differentiation.

**Supplementary Figure 4.**
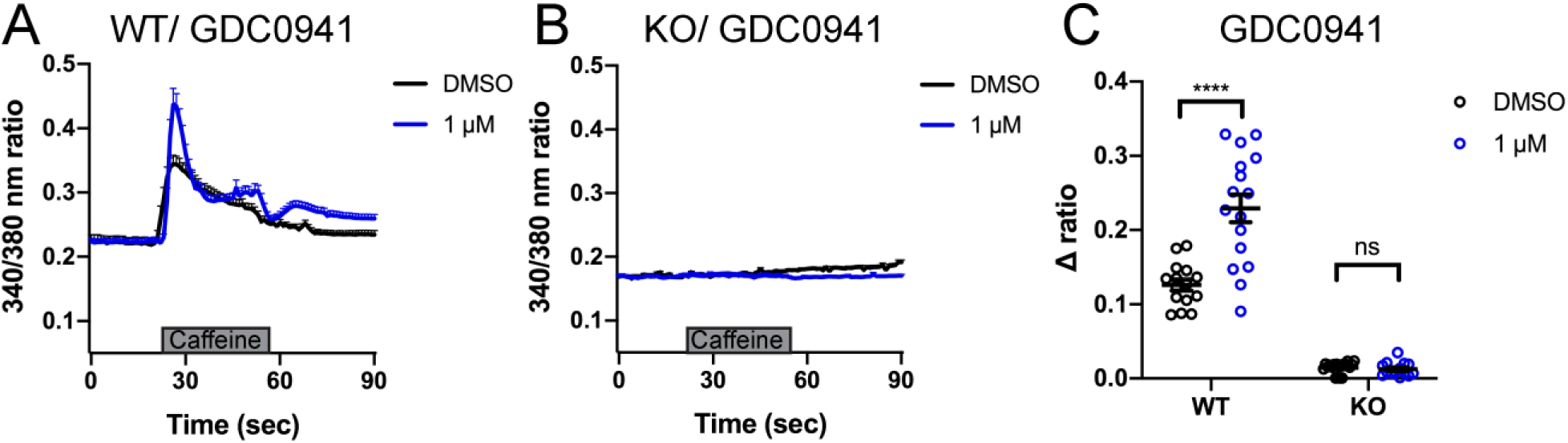
Intracellular calcium measurement in myotubes after SB203580 treatment. **A-B)** Ratiometric fura-2 imaging with 10 mM caffeine induction. **A)** WT: DMSO; n=15, 1 μM; n=16. **B)** KO: DMSO; n=19, 1 μM; n=14. Data are presented as mean ± SEM. **C)** Peak changes in intracellular Ca^2+^ expressed as Δratio (Ratio peak – Ratio baseline). WT: DMSO vs 1 μM; \*\*\*\**p* < 0.0001, KO: DMSO vs 10 μM; p = 0.988. Statistical analysis by two-way ANOVA followed by Sidak’s multiple comparisons post-test.

**Supplementary Table 1.** List of 74 chemicals that suppressed the synthetic, nemadipine-induced *unc-68* growth arrest.

## Other Supplementary Data

Results of testing other “hits” on zebrafish *ryr1b* mutants:

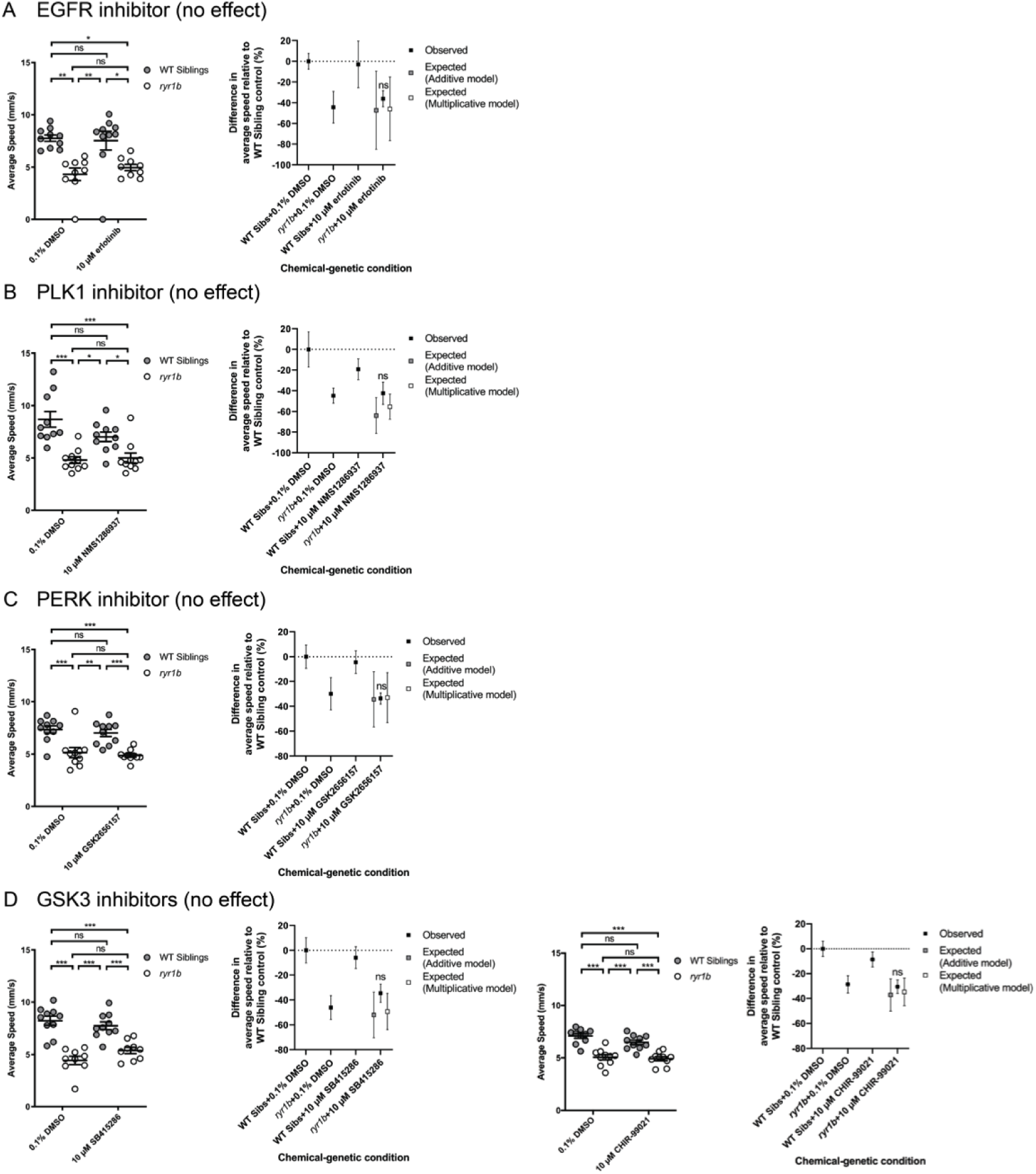
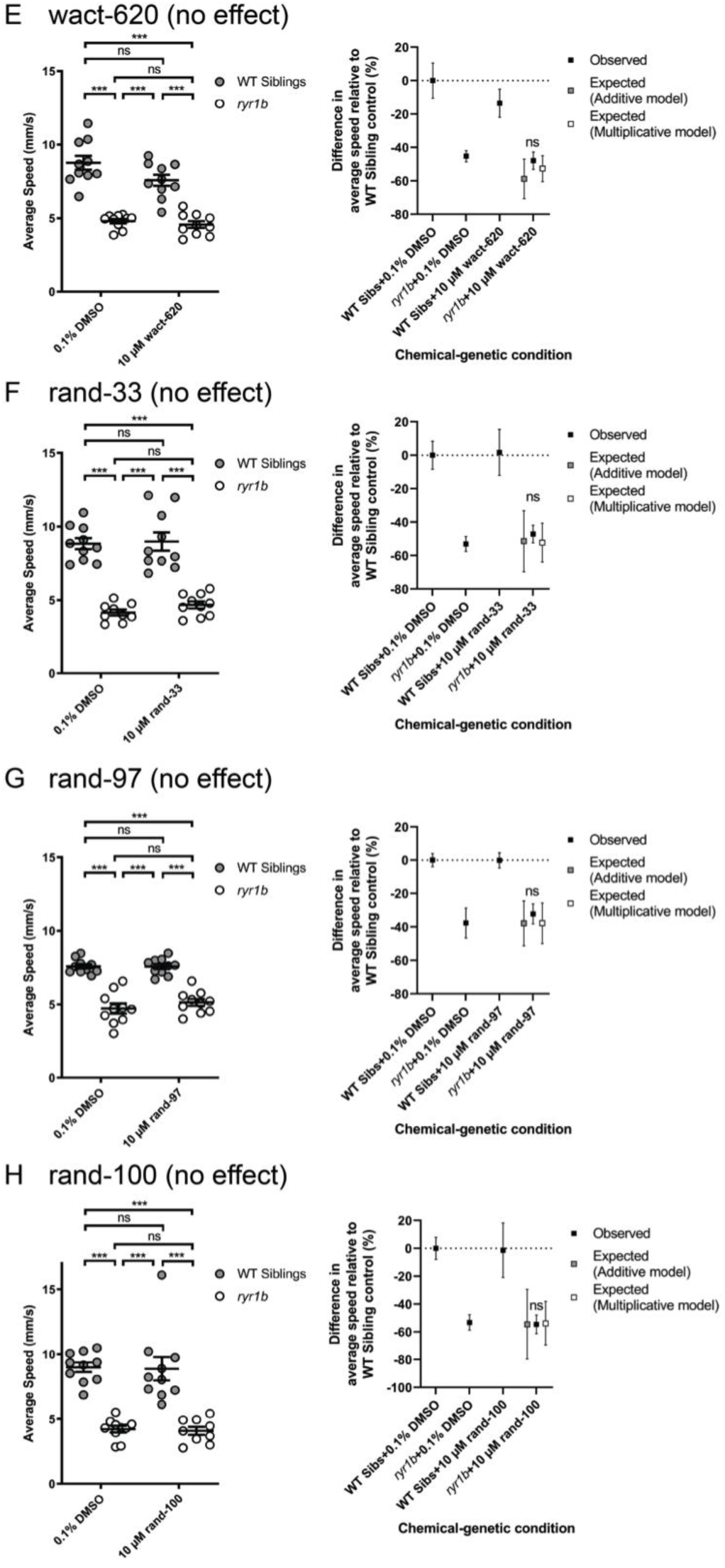
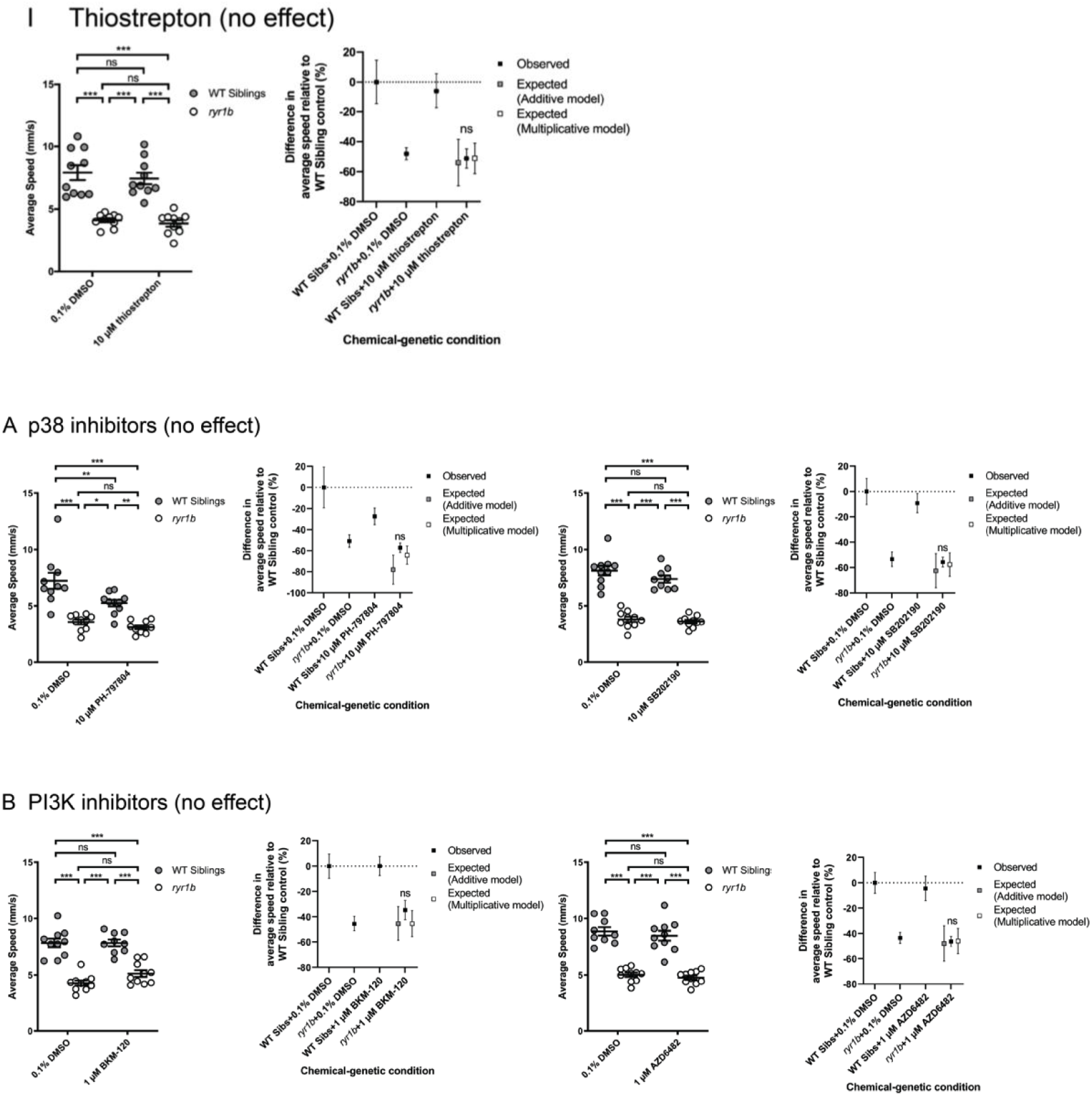

